# Mutation rates and the evolution of germline structure

**DOI:** 10.1101/034298

**Authors:** Aylwyn Scally

## Abstract

Genome sequencing studies of *de novo* mutations in humans have revealed surprising incongruities with our understanding of human germline mutation. In particular, the mutation rate observed in modern humans is substantially lower than that estimated from calibration against the fossil record, and the paternal age effect in mutations transmitted to offspring is much weaker than expected from our longstanding model of spermatogenesis. I consider possible explanations for these discrepancies, including evolutionary changes in life history parameters such as generation time and the age of puberty, a possible contribution from undetected post-zygotic mutations early in embryo development, and changes in cellular mutation processes at different stages of the germline. I suggest a revised model of stem cell state transitions during spermatogenesis, in which ‘dark’ gonial stem cells play a more active role than hitherto envisaged, with a long cycle time undetected in experimental observations. More generally I argue that the mutation rate and its evolution depend intimately on the structure of the germline in humans and other primates.

## The germline mutation rate

All evolutionary processes depend on the flow of genetic information from one generation to the next, and as with any signal, errors in transmission can occur. The rate at which this happens is called the germline mutation rate, and is of central importance to evolutionary genetics. Not only is it key to interpreting genomic differences between individuals and populations, it also determines the timescale by which we can relate genetic data to other evidence for the evolutionary past. This relationship is not straightforward however, because although in evolutionary genetic theory the mutation rate often plays the role of a fundamental constant, in truth it evolves like any other trait, and can differ by orders of magnitude between species (1).

Estimates of the mutation rate in humans have varied according to the data and methods available. The first were made even before the nature of the DNA molecule had been established (2,3), and so were indirect and restricted to mutations causing phenotypic differences, such as at dominant disease loci. Subsequent estimates were based on phylogenetic comparisons between species, with divergence times drawn from fossil evidence. More recently, developments in genome sequencing technology have enabled mutation rate estimates based on counting *de novo* mutations, comparing closely related individuals in parent-offspring trios or larger pedigrees (reviewed in (4,5)).

In principle, phylogenetic and *de novo* estimates represent different aspects of the same approach, counting genetic differences accumulated over a number of generations. For evolutionary analyses a *de novo* estimate seems at first glance more attractive because it avoids the circularity implicit in phylogenetic calibration, particularly when comparing genetic data against fossil dates. However, the first such estimates in human trios yielded a value of 0.5 × 10^−9^ bp^−1^ yr^−1^ for single-nucleotide mutations, almost half the established phylogenetic rate, and thus implying a substantial lengthening of the evolutionary timescale if applied across all hominoid lineages (4). While such a revision may be warranted in places, particularly for recent events within the genus *Homo* and the speciation of the African great apes, a longer timescale for older events is difficult to reconcile with the primate fossil record. For example, with this rate the 4.7% sequence divergence between apes and old-world monkeys (6) implies a genetic divergence time 47 Myr ago, and hence speciation ~40 Myr ago (assuming a reasonably large ancestral population), whereas the fossil record seems consistent with a divergence no more than 25-30 Myr ago (7).

Several explanations for this disagreement have been proposed, including the possibility that *de novo* estimates have failed to correctly quantify false positives or inaccessible regions of the genome (5). However, while there are caveats to any approach, more than a dozen subsequent *de novo* studies have consistently produced similarly low values (5). This includes one study based on more distantly related individuals (8), and while other forthcoming pedigree-based estimates may lead to some adjustment (for reasons discussed below), it seems unlikely that methodological considerations alone will close the gap between phylogenetic and *de novo*-estimated rates. Furthermore, additional evidence supporting a low germline mutation rate in modern humans comes from comparisons of ancient and modern DNA (9), and a lower rate is arguably more compatible with archaeological evidence for the timing of recent events such as the divergence of Native American and East Asian populations (10).

This paper explores three alternative explanations for the rate discrepancy and discusses factors underlying the germline mutation rate which may have led to its evolution on shorter or longer timescales. Firstly, I discuss the possibility that mutation rates may have slowed due to life history changes during the last 20 million years of hominoid evolution. Secondly, I consider whether aspects of the cellular genealogy of the germline might have led to a substantial number of mutations going undetected in trio sequencing experiments. Finally, I discuss how stem cell processes in spermatogenesis affect the germline mutation rate and how our model for this might be reconciled with recent experimental observations.

## Life history changes during hominoid evolution

One possible explanation for the discrepancy between mutation rate estimates is that rates themselves may have changed during hominoid evolution. Since they are observed to differ between species across large evolutionary distances, a slowdown on this scale is not implausible *a priori* (11,12). Indeed, great apes have evolved in several ways over this time, notably increasing in body mass (13). This itself leads to an explanation for the putative slowdown based on a change in generation time (defined as the average time from zygote to zygote along a genetic lineage), since life history parameters such as generation time scale with body mass across a wide range of mammal species (14,15). Consider a simplistic model where the per-generation mutation rate *μ*_gen_ is constant and the mutation rate per year *μ* scales inversely with generation time: *μ* = *μ*_gen_ / t_gen_. Then an increase in generation time by a factor of almost two could account for the necessary reduction in yearly rate from ~1 × 10^−9^ bp^−1^ yr^−1^ in the past to 0.5 × 10^−9^ bp^−1^ yr^−1^ today (Figure 1).

**Figure 1:**
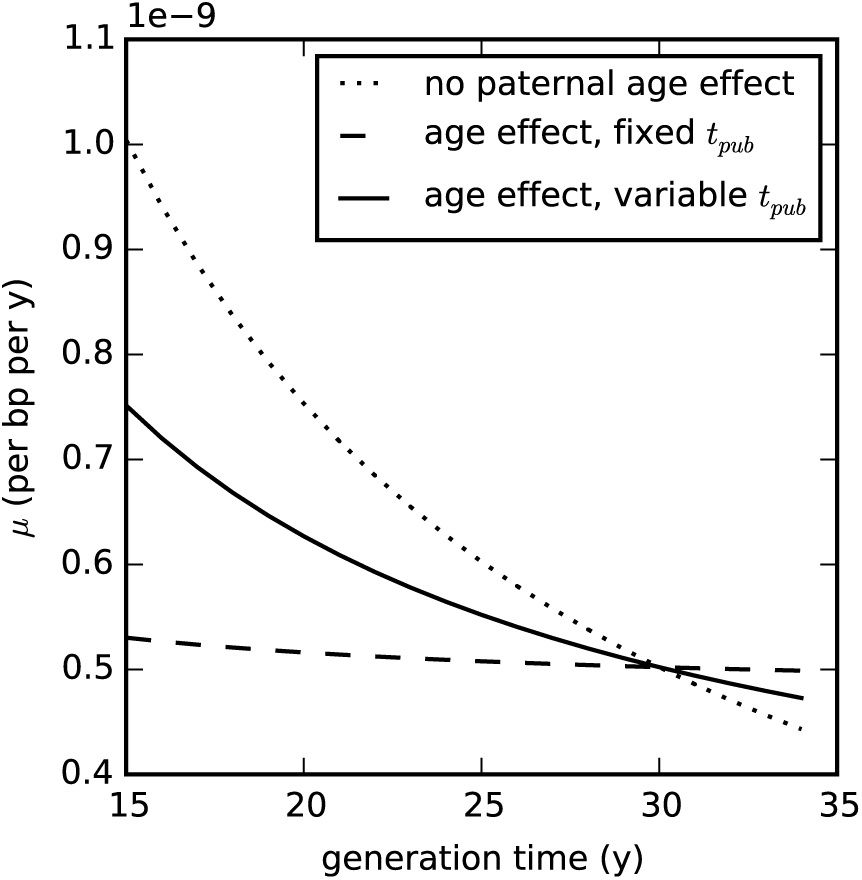
Models of human mutation rate slowdown with changing life history parameters. *Dotted*, simple scaling of mutation rate with generation time; *dashed*, including a paternal age effect but with fixed age of puberty; *solid*, including a paternal age effect and with age of puberty scaling with generation time, assuming *t*_pub_ = 14 yr when *t*_gen_ = 30 yr. Overall rates per bp are scaled to be 0.5 × 10^−9^ bp^−1^ yr^−1^ when *t*_gen_ = 30 yr.

However as previous studies have noted (5,16), this model is too simplistic, for in supposing that the number of mutations per generation is independent of *t*_gen_ it ignores the fact that older fathers tend to pass on more mutations to their offspring than younger fathers. This phenomenon, the paternal age effect, is a consequence of the fact that cell-division replication errors are the major source of germline mutation, and whereas in both sexes there are several divisions associated with embryonic development prior to gametogenesis, spermatogenesis in males involves a process of continuous further cell division throughout reproductive life. Hence the older the father, the more cell divisions his gametes will have passed through, and the more errors accumulated. By contrast in oogensis a stock of primary oocytes is generated within the developing embryo, each of which is held in stasis until the final two meiotic divisions leading to ovulation later in life.

Empirical measurements of the paternal age effect in *de novo* sequencing studies have found that the mean number of mutations passed on by fathers grows linearly with age, roughly doubling between the ages of 20 and 40 years (17–19). This would seem to largely mitigate the generation time effect on mutation rates (5,16). Consider a straightforward extension to the model presented above: for an autosomal lineage (which spends equal time in males and females) we have *μ*_gen_ = (*μ*_gen, f_ + *μ*_gen, m_) / 2, where *μ*_gen, f_ is the female mutation rate per generation and *μ*_gen, m_ the male rate. Then

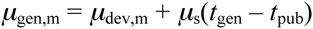

where *μ*_dev, m_ is the per-generation rate of mutations acquired during embryonic and juvenile development in males, *μ*_s_ is the yearly rate of mutation during spermatogenesis, and *t*_pub_ is the timing of puberty. We assume no age effect in mutations passed on by females, characterizing them by a single parameter *μ*_gen, f_. Parameters in this model can be taken from the experimental data of Kong *et al*. (2012) (17), who estimated whole-genome values of *μ*_gen, f_ = 14.2 and *μ*_s_ = 2.01 per yr; *μ*_dev, m_ is estimated by assuming the same mutation rate per cell division in males and females and that *μ*_gen, f_ and *μ*_dev, m_ correspond to 30 and 37 cell divisions respectively (see below for a discussion of these assumptions).

If *t*_pub_ is fixed then even a substantial change in generation time has relatively little effect on the yearly mutation rate under this model, as shown in Figure 1. However, the assumption of a fixed age of puberty is itself almost certainly invalid, since like other life history parameters the age of male sexual maturity scales with body mass across the primates, and variation within extant primates suggests a strong correlation (*R*^2^ = 0.84) with *t*_gen_ (Figure 2). Assessments of sexual maturity can vary and may not coincide with the onset of spermatogenesis in every case (20). Nevertheless if we incorporate a linear scaling of *t*_pub_ with *t*_gen_ we recover much of the generation time effect, in the sense that an increase in *t*_gen_ from 15 to 30 yr now corresponds to a reduction in *μ* by a factor of 1.5 (Figure 1).

**Figure 2.**
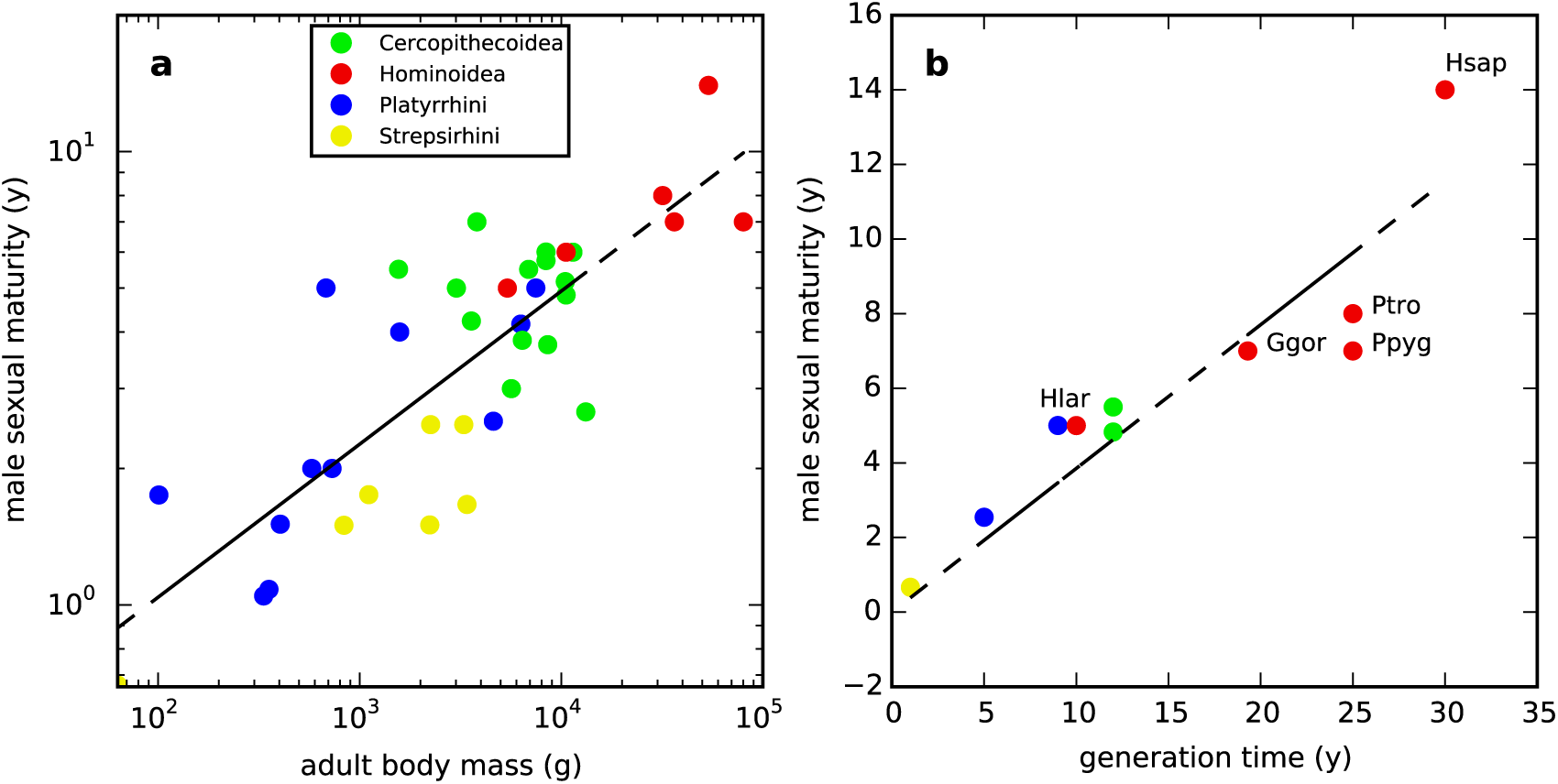
Variation of male age of sexual maturity with body mass and generation time in extant primates. (**a**) Variation with body mass suggests a scaling coefficient of 0.34. Data from (21,22). (**b**) A zero-intercept regression of age of sexual maturity on t_gen_ has slope 0.38. Data from (6,21,23–28). Hominoids are labeled: Hsap, human; Ptro, chimpanzee; Ggor, gorilla, Ppyg, orangutan; Hlar, gibbon.

## Hidden germline mutations in trio sequencing

An alternative explanation for the discrepancy in rates may lie in how *de novo* sequencing experiments relate to the cellular genealogy of the germline, and the definition of germline mutation rate as the mean number of mutations acquired on a germline lineage from zygote to zygote. Mutations on somatic lineages are important in the context of diseases such as cancer, but such lineages do not as a rule extend beyond lifespan of the organism and thus make no direct contribution to evolutionary genetic processes. However, the detection of *de novo* mutations in trios is based on sequencing somatic cells in parents and offspring, not zygotes (or even other germ cells). To understand the implications of this and how these experiments relate to what we want to measure, we need to consider the cellular genealogy of the germline within a family (Figure 3).

**Figure 3.**
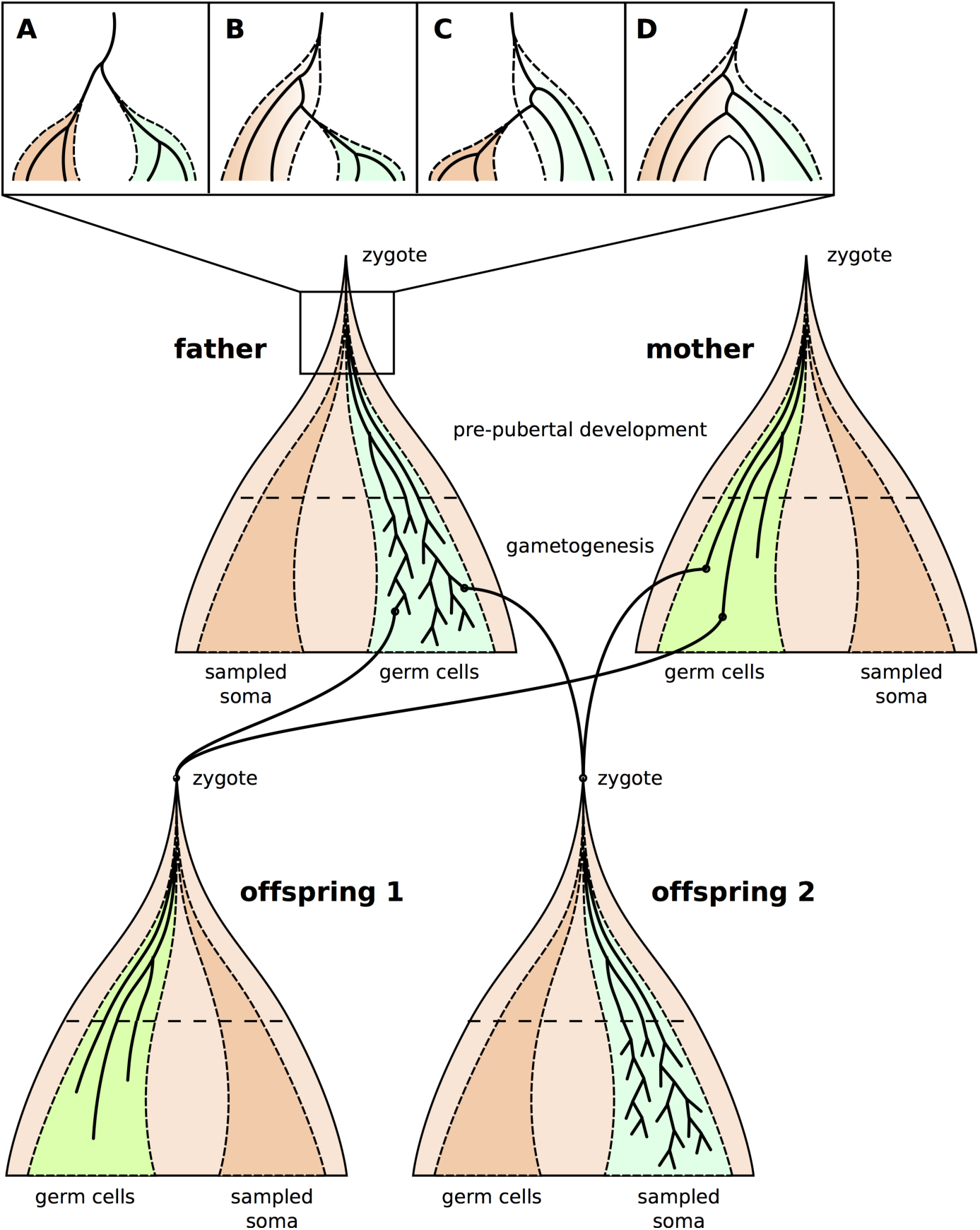
Cellular genealogies in a two-offspring family. Solid black lines represent cellular lineages; germ cell populations are shaded green (in females) or blue (in males), somatic cells are shaded orange. Darker somatic populations represent the cellular ancestors of somatic cells sampled in a sequencing experiment. Panels A-D show possible configurations of germ and sampled somatic cellular lineages at the early post-zygotic stage: A, any cell ancestral to sampled somatic and germ cells is ancestral to all such cells; B, cells may be ancestral to all germ cells but only some sampled somatic cells; C, cells may be ancestral to all sampled somatic cells but only some germ cells; D, cells may be ancestral to some germ cells and some sampled somatic cells (meaning that some germ cells may be more closely related to some somatic cells than to other germ cells, and vice versa).

Germ cell specification – the process by which certain cellular lineages are set aside as germ cells – occurs in mammals around the time of gastrulation. Following invagination of the epiblast, a number of cells originating there find a niche in the wall of the yolk sac and subsequently migrate as primordial germ cells (PGCs) to the gonadal ridge. Many somatic lineages also differentiate around the same time, and also have their origins within the epiblast. In humans this specification process occurs about two weeks after fertilization, or ~15 cell divisions (29). Thereafter, germ cell lineages undergo several further divisions in preparation for gametogenesis: approximately 15 more divisions in females and 20-24 in males (29,30). Thus in total there are about 30-40 mitotic divisions from fertilization to puberty, at which point in males the population of gonial stem cells (GSCs) is established in the testes, and the primary oocytes have been formed in females. From then on the male and female gametogenetic processes differ markedly, with GSCs replicating continuously in adult males to maintain the germ cell lineages and support gamete production.

Given this structure, the fact that *de novo* sequencing estimates are based on sequencing somatic rather than germ cells creates a potential for error. For example, in comparing parents and offspring, mutations arising early in the somatic cellular genealogy of the offspring may be counted as *de novo* germline mutations (false positive errors), and early mutations in either parent on lineages having both somatic and germline descendants may not be recognized as *de novo* (false negatives).

Some of these cases may be excluded or recovered by careful filtering based on the fraction of somatic cells in which they are present (31). However there may be a class of early post-zygotic mutations which cannot be accounted for in this way, depending on when and how the divergence of germ cell and sequenced somatic lineages occurs. Prior to the completion of this divergence, early embryonic cells may be ancestral both to germ and somatic cells within the organism, and mutations occurring then may be shared by some or all cells in either genealogy (Figure 3). Such ‘hidden’ mutations could contribute a component to the germline mutation rate which is undetectable in parent-offspring comparisons, and whose size depends on the number of cell divisions and the per-cell division mutation rate at this early stage (32). For example, it has been suggested that the first few post-zygotic cell divisions might be particularly mutagenic, based on the high level of chromosomal instability and other errors found in early IVF embryos and the frequency of early pregnancy loss after conception (33–35).

The potential for hidden mutations depends on the distribution of cell fates within the epiblast (for which much of our understanding derives from studies in mice or non-human primates). It may also depend to some extent on which somatic cells are sequenced. For example, compared to cells sampled from multiple tissues or from blood, cells derived only from one tissue or region of the body may descend from a smaller number of lineages at any given stage in development. As a consequence, configurations A and C for the divergence of germ and somatic lineages in Figure 3 may be more likely for such cells, potentially increasing the number of early cell divisions in which mutations would be hidden. As an aside, the observation that parental mosaicism in blood is correlated with recurrence risk (36) suggests that lineage ancestries for these cells at least are mixed in humans (case D in Figure 3) (37). Lineage tracing experiments on mouse oocytes suggest that a similar situation exists in mice across a range of somatic cell types, notwithstanding a degree of lineage clustering by cell type (38).

Might a hidden mutation component explain the discrepancy between phylogenetic and *de novo* rate estimates? Various considerations suggest that this is unlikely, subject to further data. Firstly, although hidden mutations are impossible to detect in single-generation experiments, comparisons over many generations should be sensitive to mutations on all ‘internal’ segments, including all hidden mutations except at the root and leaves of the pedigree. If hidden mutations make a substantial contribution to the germline mutation rate, we might expect pedigree-based estimates to be higher than those made in trios. Two such studies have been published to date, of which one did not differ significantly from trio-based estimates (8) and the other obtained a value 33% higher (39). Forthcoming studies may clarify this picture.

Secondly, a large hidden contribution should lead to a correspondingly high rate of within-family recurrence of genetic diseases associated with *de novo* mutations. This would be in addition to rates of recurrence due to shared gametic ancestry following germ cell specification, for which previous models have estimated recurrence rates of ≪1% for mutations of paternal origin (i.e. most mutations) and ~4% for those of maternal origin (36,40,41). By definition, hidden mutations occurring in a parent will be present in all of his or her gametes. Thus if hidden mutations constitute a fraction *ϕ* of all germline mutations, the probability of recurrence due to such mutations will be *ϕR*, where *R* is sibling relatedness. If hidden mutations occurred at a rate similar to the observed *de novo* mutation rate, so that the total *de novo* rate matched the phylogenetic rate, we would expect to see at least 25% recurrence of autosomal-linked diseases (*R* = 0.5). Recurrence at sex-linked loci depends on the sexes of offspring, but even in brothers of female offspring we would expect a recurrence rate of at least 12.5% for diseases caused by *de novo* mutations on the X chromosome (*R* = 0.25).

Clinically estimated recurrence rates depend on the disorder involved and the nature of the causative mutation or mutations. Some disorders such as Duchene muscular dystrophy show recurrence rates as high as 14% (42), but estimates are generally less than 1% (43,44). However, such estimates may not necessarily reflect the recurrence rates of single-nucleotide variants (SNVs) as counted in *de novo* sequencing experiments and in phylogenetic comparisons. Even where they relate to clinical genetic data, such data often include structural mutations and chromosomal abnormalities whose origins may tend to differ from those of *de novo* SNVs. In particular, chromosomal rearrangements may be enriched for meiotic errors (45), whereas the apparent linearity of the paternal age effect suggests that germline SNVs are dominated by mitotic events. Where clinical estimates are based on phenotypic recurrence, uncertainty arises in modeling the relationship between genotype and phenotype, the number of loci involved and controlling for possible environmental factors. Additionally, some phenotypes may be difficult to diagnose consistently, particularly where there is already a diagnosis in siblings, and further potential bias arises from stoppage, whereby parents of an affected child are less likely to have additional children (46,47). Some of these considerations suggest that clinical estimates might underestimate the true recurrence of *de novo* germline mutations. However, the degree of underestimation would have to be at least an order of magnitude to be consistent with a substantial contribution of hidden mutations to the germline mutation rate.

Another effect of hidden mutations would be to inflate the male-female mutation rate ratio as measured in trio comparisons. If hidden mutations occur with equal probability in males and females, and if the male-female ratio of observed (non-hidden) mutations is *α*_obs_, it is straightforward to show that the true male-female ratio *α* is bounded above by *α*_obs_ and given by

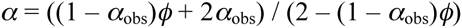

The value *α*_obs_ estimated in trios by Kong *et al*. (2012) (17) was 3.9. Alternative estimates based on comparing X-chromosome and autosomal genetic diversity within or between species (which should reflect all germline mutations) have typically fallen in the range 3–7 (48). Values at the lower end of this range could suggest a moderate contribution from hidden mutations (rearranging the above expression, *α* = 3 and *α*_obs_ = 3.9 implies *ϕ* = 0.15), but a value higher than *α*_obs_ would be inconsistent with the model presented here. In this case it may be that hidden mutations are not equally likely in males and females, which would be surprising given that most or all of the cell divisions involved occur prior to the onset of somatic sex differentiation in the embryo. However there are also several factors which can substantially bias the rate estimated from X-autosomal comparisons, including selection, sexbiased demography and differences between male and female generation times (49–51). Moreover, just as the overall germline mutation rate may have varied over evolutionary timescales, so too might the gametogenetic factors contributing to male mutation bias, which would further affect estimates based on genetic diversity.

## Changes in the structure of spermatogenesis

The importance of paternally transmitted mutations focuses attention on spermatogenesis as a key factor affecting the germline mutation rate. The established model of human spermatogenesis is based on long-standing experimental observations of the seminiferous epithelium (the environment within the testes where spermatogenesis occurs) (52,53). Yves Clermont observed several types of spermatogonial cell in humans, differing in their appearance and degree of staining with hematoxylin and eosin (54). Two of these types correspond to self-renewing (GSC) states (55), and based on their staining are generally referred to as pale (A_p_) and dark (A_d_) spermatogonia. However in Clermont’s observations only A_p_ cells were seen to actively divide, each doing so every 16 days to produce a new A_p_ cell and a progenitor spermatocyte which he termed B-spermatogonia. The latter subsequently undergo two further mitotic divisions and meiosis to produce up to 32 spermatozoa (55) in a process lasting 48 days. Thus A_p_ cells are widely regarded as the active spermatogenetic population, and A_d_ cells are thought to comprise a pool of reserve stem cells, to be drawn upon only when the active population has failed or is damaged. Figure 4 (model 1) illustrates this model in terms of the cell states and transitions involved.

**Figure 4.**
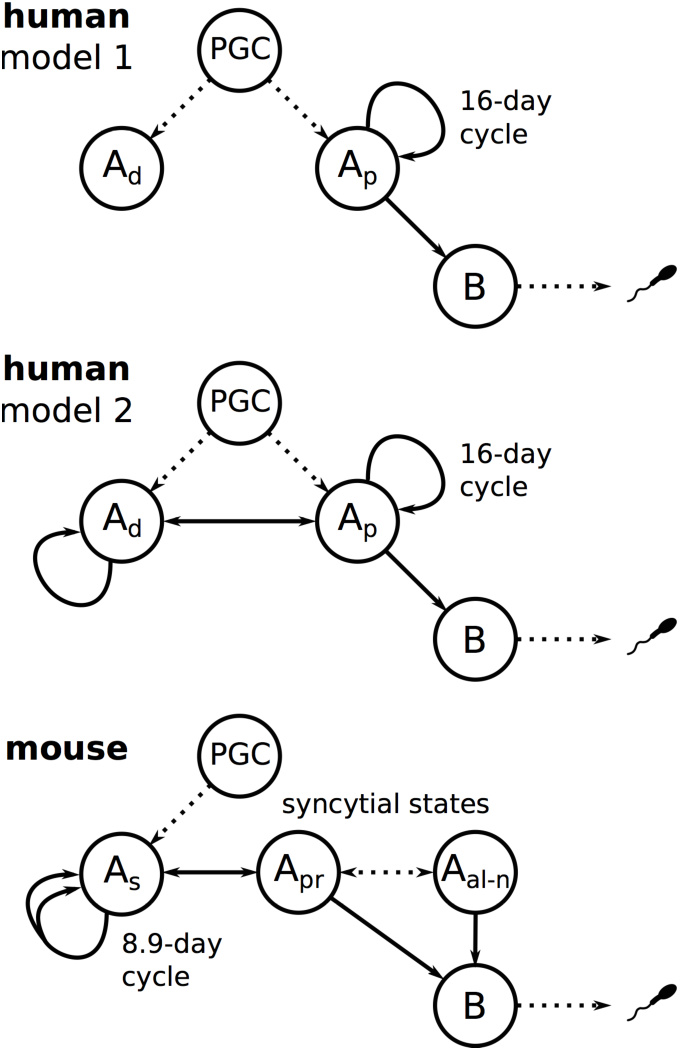
Cell states and possible transitions in models of human and mouse spermatogenesis, from primordial germ cell (PGC) to spermiogenesis. Dotted lines represent transitions involving one or more non-replicating intermediate states (for example, the initial transition from PGC to A_p_ is sometimes thought to pass through a temporary A_d_ state). Two human models are shown: model 1 is the established model originally due to Clermont (52), and model 2 an alternative model discussed in the text. In the mouse model, the double arrow from A_s_ to itself indicates that cell division is symmetric: A_s_ → A_s_ + A_s_.

One possible explanation for a slowdown in mutation rate would therefore be an increase in the cycle time of the seminiferous epithelium during hominoid evolution, leading to fewer mutations acquired during spermatogenesis for a typical adult male. Such a change is equivalent to varying *μ*_s_ in the model discussed above, and Figure 5 shows the effect on germline mutation rate, assuming that puberty also scales with generation time as previously discussed. The seminiferous epithelial cycle time in monkeys varies between 9 and 11 days (56), and if the cycle time in ancestral great apes was similar to this, the change since then would correspond to a mutation rate slowdown by almost a factor of two (dashed line in Figure 5), and perhaps sufficient to explain the discrepancy in mutation rates.

**Figure 5.**
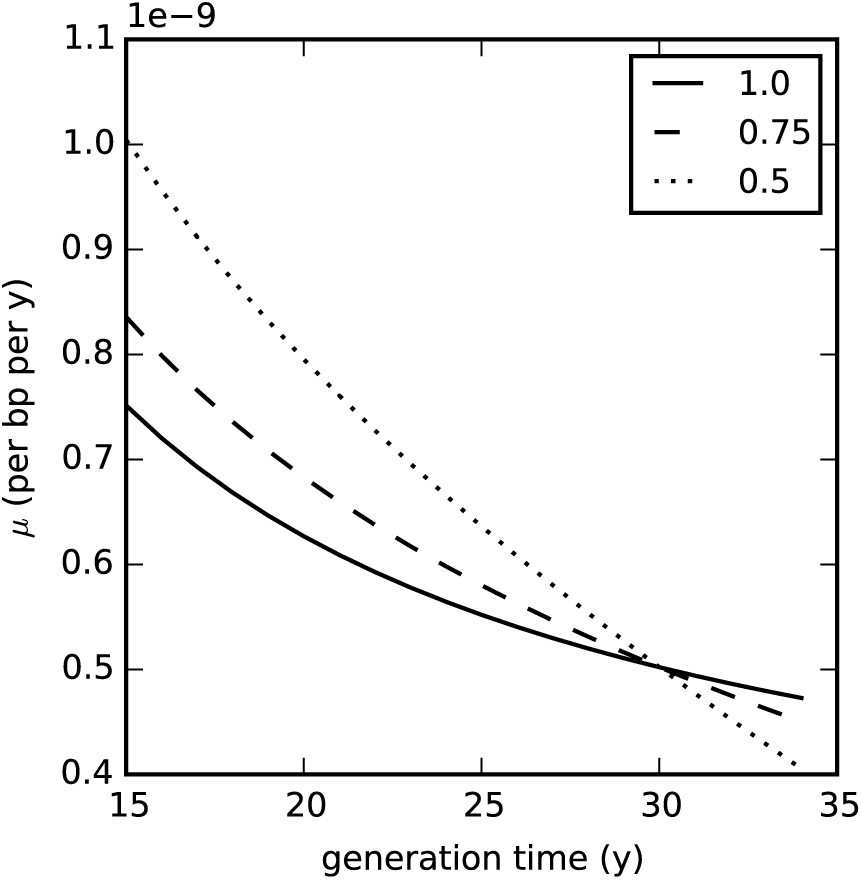
Models of human mutation rate slowdown in which the seminiferous epithelial cycle time increases with generation time and age of puberty. *Dotted line*, cycle time is 0.5 times the present length when generation time is 15 y; *dashed line*, cycle time is 0.75 times the present length when generation time is 15 y; *solid line*, no change in cycle time (identical to the solid line in Figure 1). Overall rates per bp are scaled to be 0.5 × 10^−9^ bp^−1^ y^−1^ when *t*_gen_ = 30 y.

However there is a problem with this model as presented, in that lineages resulting in gametes produced by a 30-year-old male would have passed through ~400 cell divisions since fertilization, meaning we would expect a roughly tenfold increase in the number of mutations passed on to offspring at age 30 compared to those at puberty. Yet sequencing studies have consistently measured only a twofold increase from early to late adulthood (17–19). This discrepancy, noted also by Ségurel *et al*. (5), suggests that aspects of the model for spermatogenesis need to be revised.

One possibility is that per-cell-division mutation rates are much higher at earlier developmental stages than during spermatogenesis. For example, there is evidence that somatic cell division mutation rates are substantially higher than those in germ cells (57) and it may be that changes in the environment or phenotype of cells at germ cell specification are accompanied by improved mechanisms of DNA replication error correction. In order to account for a weak paternal age effect, the mutation rate in the first 15 cell divisions (prior to specification) would need to be a factor of ~20 higher than in subsequent divisions, which is near the limit of the range of reported estimates for germline and somatic cells (1). Note also that this excludes any hidden mutations of the kind discussed above, although such mutations would also be made more likely by an elevated post-zygotic mutation rate. Alternatively, elevated per-cell-division rates may last for a longer time, perhaps until the onset of spermatogenesis or shortly thereafter. If we assume a higher rate applies to the first 40 divisions then it need only be higher by a factor of ~9. A recent study of transmitted mutations in a large cohort including some teenage fathers suggested that the very earliest stage of spermatogenesis may be more mutagenic (19). If true, this might reflect a shift to lower mutation rates once spermatogenesis is fully established. Early spermatogenesis is known to differ in some respects from the process later on: for example daily sperm production volumes are ~10 times lower in teenage males than in men 20-30 years old (58).

Another possibility is that the apparent 16-day cycle of the seminiferous epithelium is only part of the picture and that germline lineages actually experience a longer cell cycle time for some or all of their passage through spermatogenesis. This would imply a more complex structure for GSC state transitions and the number of self-renewing states in which they can exist. It is of course possible to conceive of many such models, but a relatively straightforward extension of the existing model would be for the A_d_ cells to play a more prominent role. If they replicate with a longer cycle time than was detectable in Clermont’s data, and if transitions are possible between the A_d_ and A_p_ states, then germline lineages could spend some or even most of their time in the A_d_ state during spermatogenesis (Figure 4, model 2). By reducing the number of germline cell divisions, this could account for a weaker than expected paternal age effect.

Potential evidence for such a model comes from several sources. Within primates, investigations of spermatogonial renewal in monkeys after exposure to radioactive or contraceptive agents (59,60) have shown that A_p_ cells may be able to transition to A_d_ without undergoing cell division. If this occurs under normal conditions then the A_d_ state could play a role other than that of a dormant and nonproliferative reserve. Other evidence comes from comparison with spermatogenesis in mice, which although differing in several respects does share many basic features with primates (61) (Figure 4). GSCs in mice can exist in a singular state (A_s_) or in various syncytial states wherein the nuclei share a common cytoplasm, either paired (state A_pr_) or in longer alignments of *n* cells (states A_al-*n*_) (62). Recent experimental studies suggest that progenitor spermatocytes may be produced from divisions of any of these states (61,63), but the degree of commitment to (or likelihood of) differentiation may be greater in the A_al_ state. An analogy can be drawn with the process in humans, based both on function and on expression levels of several molecular markers, in which the As state corresponds to A_d_ and the A_pr_ and A_al_ states correspond to A_p_, (61,64). Moreover, a model of stochastic transition between the A_s_, A_pr_ and A_al_ states, in which intracellular bridges are broken as well as created, has been shown to fit the dynamics of GSC populations in mice (63,64). The analogous set of transitions in humans and other primates would fit the alternative model shown in Figure 4.

By making certain assumptions about the possible transitions between GSC states, we can estimate what cycle time for the A_d_ state would produce the observed paternal age effect. Previous approaches to modelling stem cell systems have sometimes assumed homeostatic equilibrium as a way of estimating or constraining model parameters (e.g. (63)), but it may be that spermatogenesis is better represented as a near-equilibrium process. For example there is notable variation in daily sperm production (DSP) with age: in humans mean DSP decreases steadily in older men, dropping by a factor of two from age 20 to 80 years (58). To capture these non-equilibrium aspects we can use a finite state simulation in which transitions between cell states have both a probability parameter and an associated transition time (Figure 6). The basic assumption of this model is that the transition A_p_ → A_d_ occurs with some probability during each A_p_ cycle. For the reverse transition, since A_p_ is observed to divide asymmetrically (A_p_ → A_p_ + S) I assume for now that A_d_ behaves similarly (A_d_ → A_d_ + A_p_); an alternative choice (A_d_ → A_p_) is discussed below. I also introduce a cell death state which both regulates the process (since otherwise A_d_ replication will lead to unbounded GSC proliferation) and ensures that gamete production declines with age. Cell death likely plays a role in regulating many stem cell systems (65), and in spermatogenesis GSC replication occurs only within a niche at the basement membrane of a seminiferous tubule for which several cellular and other environmental factors may be essential. In particular the availability of Sertoli cells, somatic cells which play both a structural and regulatory role in gamete production, is thought to be a critical factor (53,58). Cell death probabilities in this model are an abstraction representing the typical availability of and contention for these critical factors.

**Figure 6.**
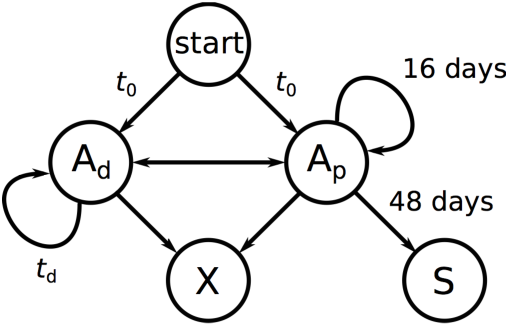
A possible finite state model for spermatogenesis. Germline lineages are traced through the model, entering at the START state, from which they can transition to either of the GSC states A_d_ or A_p_. The cell death state (X) can be reached from either GSC state, whereas we assume here that spermiogenesis (S) is possible only from A_p_. Transitions from the GSC states correspond to cell divisions, so the model describes a branching process representing an entire genealogy of spermatozoa, each descended from a single primordial germ cell (PGC) and having an associated age and total number of cell divisions (depth) from the root of the genealogy. In the established model (model 1 in Figure 4), the transition A_p_ → A_d_ has zero probability under normal conditions.

A simple parameter-space search, fitting simulated output of this model to the observed slopes of the paternal age effect and DSP age profile, estimates an A_d_ cycle time of around 300 days and values of 20-30% for transition probabilities to the cell death state (Figure 7). Replication of A_d_ cells on this timescale would likely not have been observed in Clermont’s or subsequent experiments, although transitions A_p_ → A_d_, which in this example are predicted to occur in 10% of A_p_ cycles, might be detectable.

**Figure 7.**
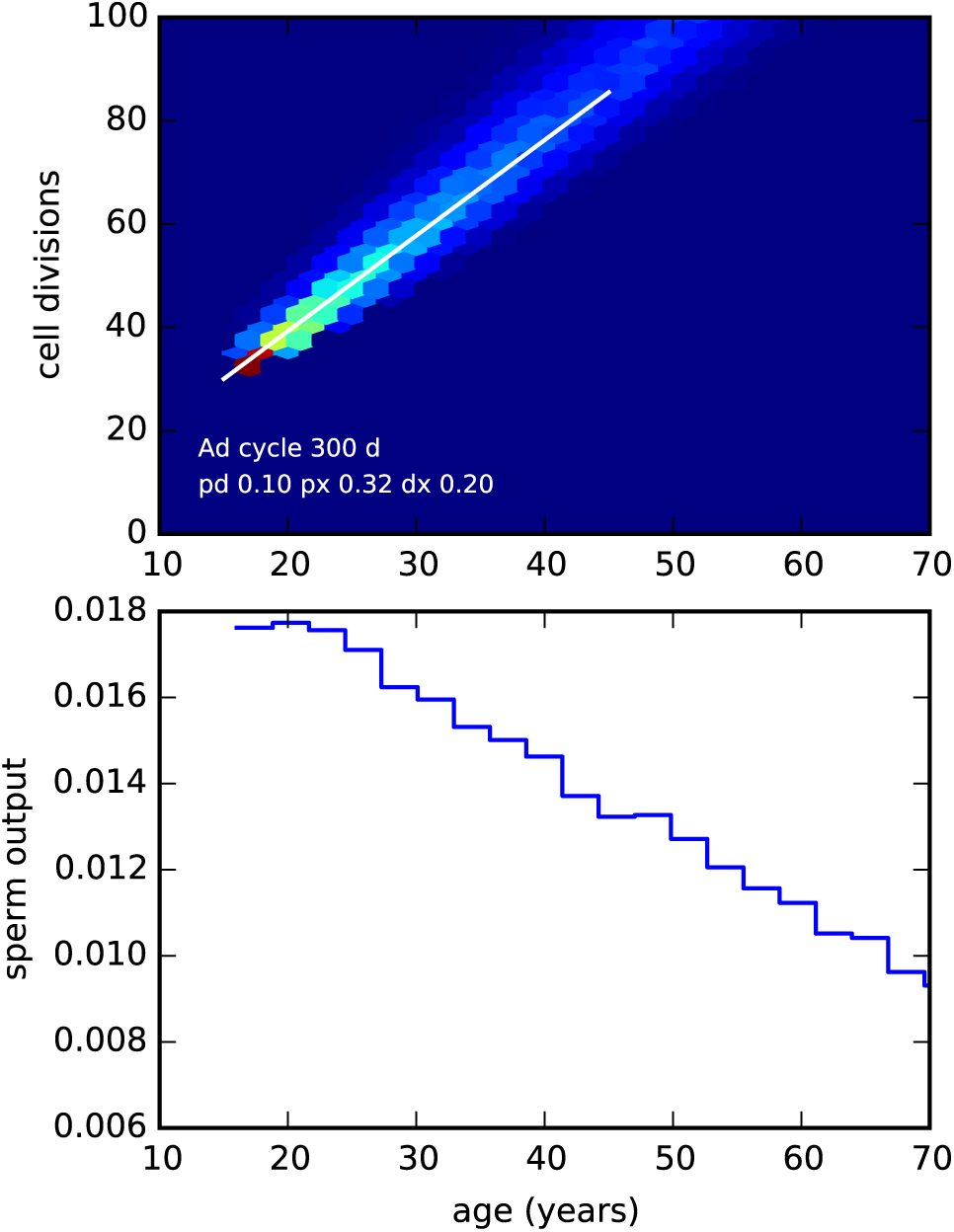
Output from an implementation of the model in Figure 6, for parameters fitting the observed paternal age effect and decline in sperm production. (Code available on request.) *Upper panel*: Joint distribution of paternal age and number of cell divisions since the zygote in simulated spermatozoa (colours indicate relative frequency from zero (dark blue) to high (red)) for a set of parameters in which the A_d_ cycle time *t*_d_ = 300 days and initial transition times *t*_0_ were sampled uniformly from the range 5000–5500 days, P(A_p_ → A_d_) = 0.1, P(A_p_ → X) = 0.32, P(A_d_ → X) = 0.2. The white line shows the slope of the paternal age effect observed in humans. *Lower panel*: Simulated relative daily sperm output as a function of age.

Other model assumptions are possible, and may result in different parameter estimates. For example, in a model where the A_d_ → A_p_ transition occurs without cell division, so that the A_d_ cycle is essentially a quiescent interlude before GSCs return to the active A_p_ state, a similar procedure fitting the observed paternal age effect estimates a cycle time of around 750 days (data not shown). However the point here is not that parameters can be inferred under one model or another, but the fact that including an active role for the A_d_ cells allows models which are compatible both with longstanding observations of the seminiferous epithelium and with recent measurements of the paternal age effect. More sophisticated models might also include feedback or global regulation mechanisms other than cell death (53,56), age-related changes in cell-division mutation rate and spermiogenetic efficiency, and perhaps phenomena such as selfish spermatogonial selection (66). At present experimental data is limited, but more extensive data including trio sequencing on population-wide scales will provide a better basis for exploration of spermatogenetic models along these or similar lines.

## Discussion

I have argued that the discrepancy between phylogenetic and *de novo* estimates of the mutation rate is more likely due to a genuine evolutionary slowdown than to methodological errors or the failure of trio sequencing experiments to detect early post-zygotic mutations. Nevertheless, the latter factors may be present at some level and thus contribute to the discrepancy, meaning that the magnitude of any slowdown may be less than was first hypothesized. Also, while we may be confident that rates have slowed at some point during primate evolution, our inference of the timing, extent and number of ancestral lineages involved in such a slowdown is determined largely by the fossil record and the confidence with which we can constrain speciation events, particularly within the hominoids. Initial attempts to reconcile the rate discrepancy were concerned with the plausibility of a slowdown affecting all four great ape lineages in parallel and to the same degree, given that their branch lengths from an outgroup such as macaque do not differ substantially (50). However, if newer interpretations of the fossil record were to admit a more ancient speciation time of 20 Mya or more between the ancestors of orangutans and other great apes, they would be consistent with an earlier slowdown affecting only the stem hominoid lineage, and this would suffice to explain the current data without requiring parallel evolution.

More broadly, and regardless of the extent to which rates may have changed in recent primate evolution, the processes considered here are relevant to evolutionary genetic analyses across the mammalian tree and beyond. Previous studies have proposed life history variation as an explanation for mutation rate change, but it is clear that such explanations need to involve more biologically sophisticated models incorporating factors such as varying pubertal age and sex-dependent parameters (5,51). Mutation rate change may also be due to evolution in the underlying cellular processes and genealogical structure of the germline, particularly in gametogenesis. Here too, recent experimental data are incongruous with existing models of spermatogenesis and the strength of the paternal age effect. I have focused on potential variation in cell-division mutation rates at different developmental stages and on the stem cell states involved in spermatogenesis. Other issues not touched on include the relative importance of spontaneous mutation processes (67), potential evidence for a maternal age effect (68), and the evolution of regulatory factors controlling gametogenesis (53). Progress to date in addressing these questions has been difficult in part because of the challenge of obtaining experimental data on human germline processes: some techniques can only be applied to non-human models, and genome sequence data for human *de novo* mutations has previously been limited. However the potential now exists for large-scale genome sequencing of somatic and germ cells and experimental studies of non-human and human stem cell systems. In combination with computational modeling approaches such as that presented here (and widely used in previous studies to explore stem cell population dynamics (36,63,69–71)), these developments will facilitate a better understanding of mutation processes and the evolution of the human germline.

